# Spinocerebellar Ataxia Type 1 protein Ataxin-1 is signalled to DNA damage by Ataxia Telangiectasia Mutated kinase

**DOI:** 10.1101/701953

**Authors:** Celeste E. Suart, Alma M. Perez, Ismael Al-Ramahi, Tamara Maiuri, Juan Botas, Ray Truant

## Abstract

Spinocerebellar Ataxia Type 1 (SCA1) is an autosomal dominant neurodegenerative disorder caused by a polyglutamine expansion in the ataxin-1 protein. Recent genetic correlational studies have implicated DNA damage repair pathways in modifying the age at onset of disease symptoms in SCA1 and Huntington’s Disease, another polyglutamine expansion disease. We demonstrate that both endogenous and transfected ataxin-1 localizes to sites of DNA damage, which is impaired by polyglutamine expansion. This response is dependent on ataxia telangiectasia mutated (ATM) kinase activity. Further, we characterize an ATM phosphorylation motif within ataxin-1 at serine 188. We show reduction of the *Drosophila* ATM homolog levels in a ATXN1[82Q] *Drosophila* model through shRNA or genetic cross ameliorates motor symptoms. These findings offer a possible explanation as to why DNA repair was implicated in SCA1 pathogenesis by past studies. The similarities between the ataxin-1 and the huntingtin responses to DNA damage provide further support for a shared pathogenic mechanism for polyglutamine expansion diseases.

## INTRODUCTION

Spinocerebellar Ataxia Type 1 (SCA1) is an age-onset neurodegenerative disorder caused by expanded CAG DNA triplet repeats within *ATXN1 (1)*. ATXN1 is typically encoded with 6-42 CAG repeats, with a CAT interruption every twenty one repeats (1). Patients with 39-84 uninterrupted repeats will experience SCA1 symptoms; including ataxia, muscle weakness, and difficulty with speaking and swallowing (1). This CAG expansion encodes an expanded polyglutamine tract within the ataxin-1 protein, leading to neurodegeneration within the cerebellum and brainstem (2). SCA1 is a member of the age-onset polyglutamine expansion disease family, which includes Huntington’s Disease (HD) (3).

A hallmark of polyglutamine expansion diseases is that a greater number of repeats correlates to an earlier age at onset (AAO) of symptoms (4). This correlation has been well established in SCA1, however, only 70% of the variance in AAO can be attributed to CAG length (4,5). This implies that other genetic or environmental factors affect the progression of SCA1. Single-nucleotide polymorphism (SNP) analysis conducted on a cohort of HD and SCA patients found that variations in DNA repair genes significantly modified AAO for SCA1 patients (6). This was a follow-up to the landmark genetic modifiers of Huntington’s disease genome-wide association study (GWAS), which also found a significant association between DNA repair pathways and AAO in HD patients (7). The association of DNA repair pathways and progression of both SCAs and HD suggests there may be a common pathogenic mechanism for polyglutamine expansion diseases.

We have previously characterized the response of the HD protein huntingtin to DNA damage (8). Huntingtin localizes to sites of DNA damage caused by irradiation and oxidative stress (8). This response is dependent on ataxia-telangiectasia mutated (ATM), a serine/threonine kinase activated by oxidative stress, chromatin reorganization, and DNA breaks (9–11). Oxidative stress, induced by reactive oxygen species, has been suggested to play a role in multiple forms of neurodegeneration, including HD, Alzheimer’s Disease, and Parkinson’s Disease (12–14). Level of oxidative stress has also been associated with disease severity in the related disease SCA7(15).

The huntingtin response to DNA damage (8) genetic studies indicating HD and SCA1 may share a common pathogenic mechanism involving DNA repair (6,7) led us to hypothesize that ataxin-1 may have a similar response to DNA damage. Using both endogenous and transfected protein models, we show that ataxin-1 localizes to sites of DNA damage. This localization is affected by polyglutamine expansion of the ataxin-1 protein. We demonstrate this response is dependent on ATM kinase activity and characterize an ATM substrate LSQ motif in ataxin-1 at serine 188. Further, we demonstrate in a SCA1 *Drosophila* model that reduction of ATM protein levels improves motor phenotype. These results offer a possible biological mechanism for the identification of DNA repair genes as modifiers of SCA1 progression (6) and lay groundwork for future investigation of ataxin-1 as a potential DNA repair protein.

## RESULTS

To determine if endogenous ataxin-1 can localize to sites of DNA damage, immunofluorescence was performed on hTERT-immortalized retinal pigment epithelial (RPE1) cells following irradiation of regions of interest with a 405nm laser. Post-irradiation, endogenous ataxin-1 was observed to co-localize with huntingtin at the irradiated area (Fig. 1a).

**Figure 1.**
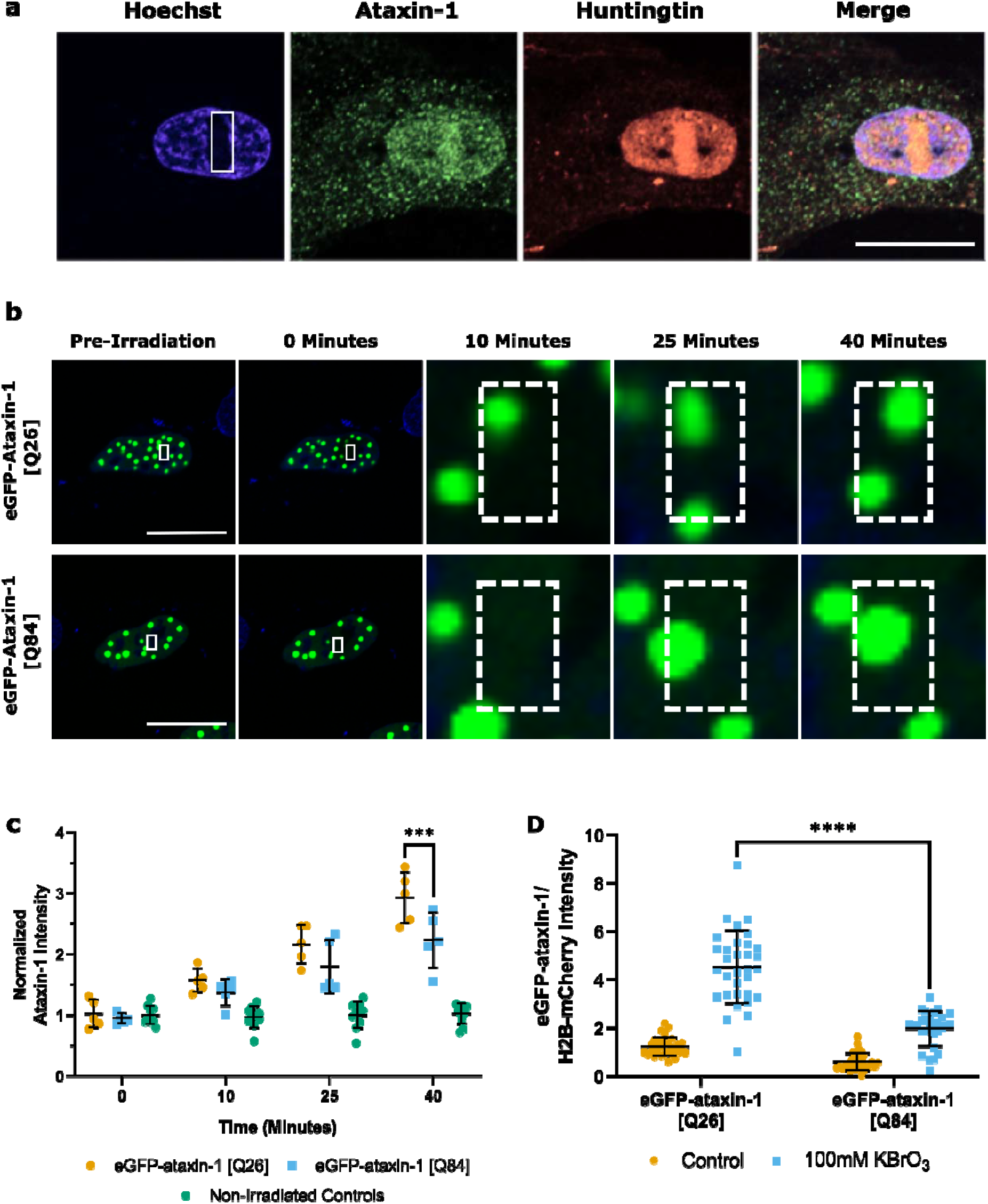
Ataxin-1 localizes to sites of DNA damage following irradiation and oxidative stress. All graphs display mean (± SD). Significance values calculated by two-way ANOVA, *** = p < 0.001, **** = p ≤ 0.0001. **a.** Laser microirradiation assay of endogenous ataxin-1 from RPE1 cell. White rectangle indicates irradiated area. **b.** Laser microirradiation assay of the indicated transfected ataxin-1 constructs in RPE1 cells. White rectangle indicates irradiated area. **c**. Quantification of part b. Average pixel intensity was calculated for control (0% laser power) and irradiated (100% laser power) regions. Values normalized to the average pre-irradiation control value to account for random movement of ataxin-1 puncta. Both constructs had significant difference from non-irradiated controls starting at 10 minutes. **d.** Chromatin retention assay with indicated transfected ataxin-1 constructs. Nuclear eGFP intensity was quantified after extraction of soluble proteins. Potassium bromate treated conditions both significantly different from controls (p ≤ 0.0001). Approximately 30 images were taken per condition with 20-40 cells per image.

Next, we asked whether increased polyglutamine tract length beyond the pathogenic threshold of 37-39 repeats would impact ataxin-1 localization to sites of DNA damage. Polyglutamine tract length above 37-39 repeats causes SCA1(16). To explore the effects of these mutations, we used an eGFP-fused full length ataxin-1 construct that we have previously characterized(17). Live cell imaging following micro-irradiation was conducted on RPE1 cells expressing each eGFP-ataxin-1 construct (Fig. 1b). Regions without ataxin-1 puncta were irradiated with an 405nm laser to induce localized DNA damage, then average pixel intensity of these regions was monitored over time. As ataxin-1 puncta localized into the irradiated region, the average pixel intensity would increase. Similar to endogenous ataxin-1, wild type ataxin-1-eGFP puncta relocated to regions of DNA damage (Fig. 1b,c). EGFP-ataxin-1 [Q84], with an expanded polyglutamine tract, had impaired relocation compared to wild type ataxin-1 [Q26] after 40 minutes (Fig. 1c). Thus, the pathogenic polyglutamine expansion affected the ability of ataxin-1 nuclear puncta to relocate to regions of damaged DNA.

We examined if ataxin-1 was recruited to chromatin following treatment with 100mM potassium bromate (KBrO_3_), an oxidizing agent which induces DNA base damage(18,19). Protein chromatin retention assays were performed on RPE1 cells transfected with eGFP-ataxin-1 constructs and histone H2B-mCherry transfection efficiency control. Both ataxin-1 constructs displayed recruitment of ataxin-1 to chromatin in response to oxidative stress (Fig. 1d). Consistent with micro-irradiation experiments, wild type eGFP-ataxin-1 had the highest level of chromatin retention, while polyglutamine expanded eGFP-ataxin-1 displayed impaired retention compared following oxidative stress (Fig. 1d). Thus, ataxin-1 chromatin association is induced by multiple types of damaging agents, and polyglutamine expansion impairs this recruitment.

We then investigated if the ataxin-1 response to DNA damage was dependent on ataxia telangiectasia mutated (ATM) kinase activity, similar to huntingtin(8). We performed microirradiation assays on RPE1 cells expressing wild type ataxin-1in the presence of 10μM KU55933, an ATM kinase inhibitor(8,20). Irradiated cells treated with ATM inhibitor exhibited a sharp decrease in overall cell fluorescence, as well as a decrease in the number of fluorescent nuclear puncta (Fig. 2a). Efflux of soluble ataxin-1 from the nucleus, followed by degredation was also observed following irradiation. This phenotype has not been previously seen in this model system (Supplemental video 1). There was no significant recruitment of ataxin-1 to sites of DNA damage in cells treated with ATM inhibitor (Fig. 2b). This suggested that ATM may directly phosphorylate ataxin-1.

**Figure 2.**
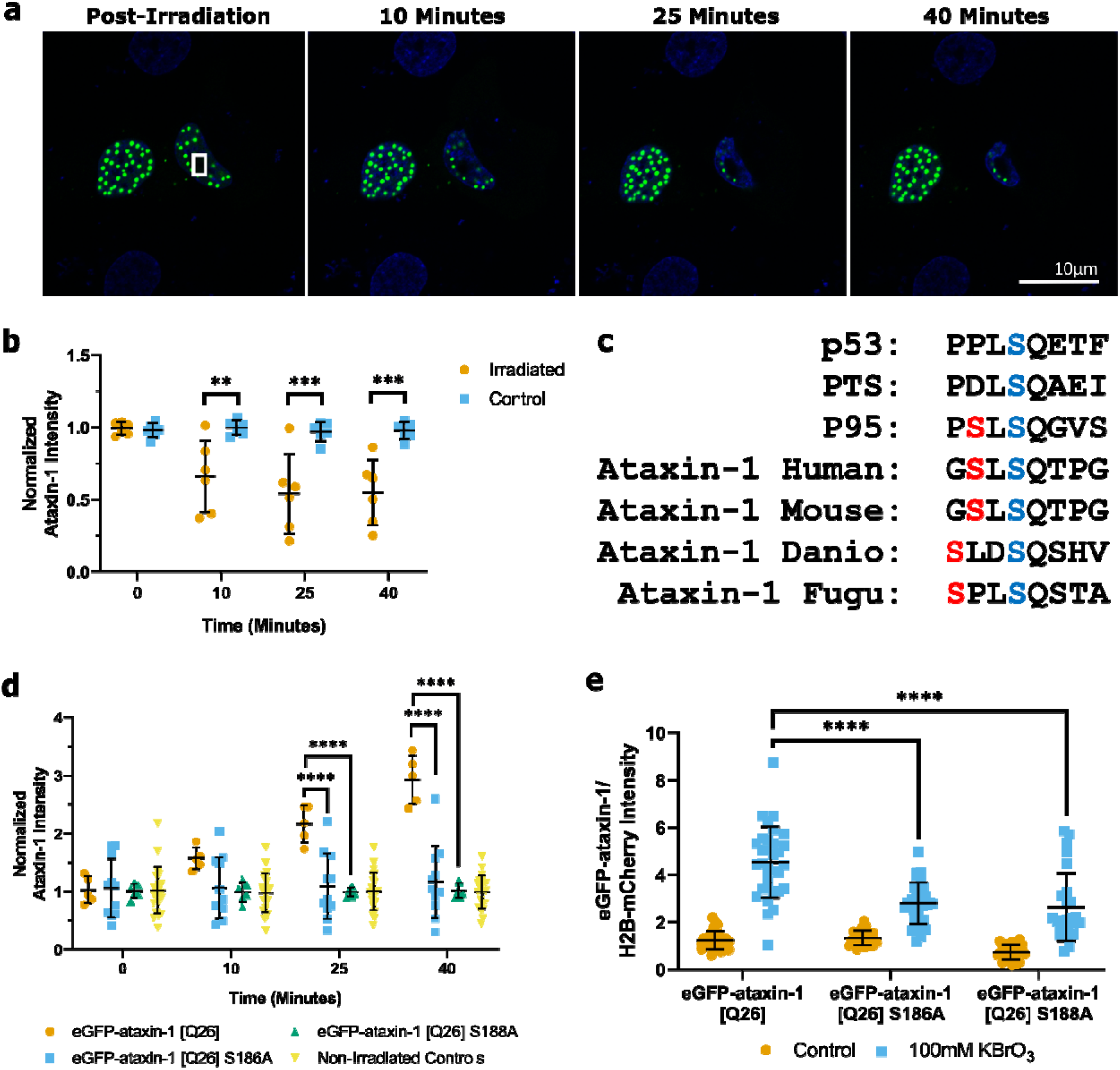
ATM inhibition prevents ataxin-1 localization to DNA damage. **a.** Laser microirradiation assay of RPE1 cells transfected with eGFP-ataxin-1[Q26] after incubation with 10μM ATM kinase inhibitor KU55933. **b.** Values from part a were quantified as in Figure 1c. Error bars show mean (± SD) on the basis of six replicates. Significance values calculated by two-way ANOVA, ** = p ≤ 0.01, *** = p ≤ 0.001. **c.** Alignment and conservation of predicted ATM kinase site in ataxin-1. p53, PTS, and p95 are canonical ATM substrate provided for comparison. Residues 185-192 of ataxin-1 shown. Adapted from Kim et al., 1999. Serine 186 shown in red, serine 188 shown in blue. **d**. Quantification of indicated eGFP-ataxin-1 construct recruitment to micro-irradiated regions. Error bars and statistical tests as described in Figure 1c. eGFP-ataxin-1 [Q26] data displayed is replicated from Figure 1c for comparison. **e.** Chromatin retention assay of eGFP-ataxin-1 [Q26] S186A and eGFP-ataxin-1 [Q26] S188A compared to eGFP-ataxin-1 [Q26], using methods and statistical analysis as described in Figure 1d. eGFP-ataxin-1 [Q26] data displayed is replicated from Figure 1d for comparison.

We used the kinase prediction software GPS 3.0 to determine if ataxin-1 contains an LSQ motif, a consensus substrate recognition motif for ATM (21,22). A putative LSQ motif was identified at serine 188 in ataxin-1. This site is conserved in ataxin-1 across species and shares homology with canonical LSQ motifs (Fig. 2c). This suggested that ATM could potentially phosphorylate serine 188 in ataxin-1. A conserved serine at position 186 was also identified in our analysis (Fig. 2c). We hypothesized it might play a priming role for serine 188, similar to ATM substrate p95 (22). Thus, we considered both serine 186 and 188 modification in our subsequent analyses.

To test whether phosphorylation of serine 186 or 188 affected ataxin-1 localization to DNA damage sites, serine to alanine substitution mutants were generated. RPE1 cells transfected with eGFP-ataxin-1 [Q26] S186A or eGFP-ataxin-1 [Q26] S188A substitutions were subjected to laser DNA damage assays (Fig. 2d). Despite similar expression levels, there was no recruitment of either alanine mutant to sites of DNA damage. In chromatin retention assays, both S186A and S188A ataxin-1 mutants had impaired recruitment to sites of damage compared to wild type ataxin-1 following oxidative stress (Fig 2e). This suggests that the two predicted ATM substrate residues are involved in regulating ataxin-1 localization to sites of DNA damage.

Based on our results, we generated an affinity purified rabbit polyclonal pS186pS188 ataxin-1 antibody. We then validated the antibody (see supplemental data).

We next examined whether modulation of ATM activity affects phosphorylation levels of endogenous ataxin-1 using the pS186pS188 antibody. Upon treatment with KBrO_3_, there was a significant increase in pS186pS188 ataxin-1 levels (Fig. 3a), indicating that this kinase substrate site was responsive to oxidative stress. Cells exposed to oxidative stress in the presence of ATM inhibitor exhibited a diminished response (Fig. 3a). Thus, oxidative stress, which increases ATM activity(23), led to increased pS186pS188 ataxin-1, while inhibition of ATM activity led to decreased pS186pS188 ataxin-1. This is consistent with DNA repair factors signaled by ATM(24).

**Figure 3.**
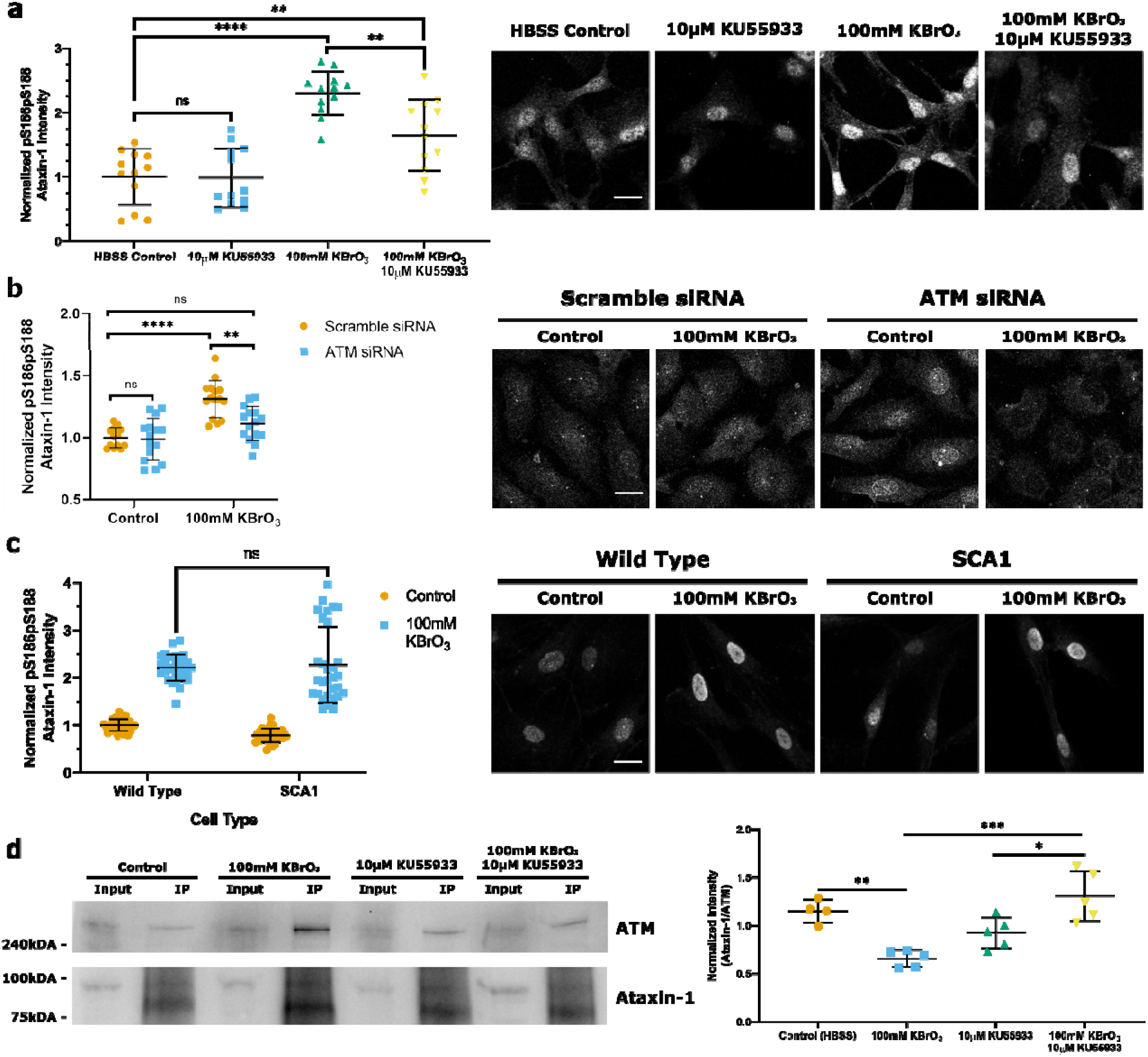
Oxidative stress increases pS186pS188 ataxin-1 levels in an ATM-dependent manner. Error bars displays SD. Significance values calculated by one-way or two-way ANOVA, * = p ≤ 0.05,** = p ≤ 0.01, *** = p 0.001,**** = p ≤ 0.0001. White scale bar indicates 10μm. **a.** RPE1 cells were treated with either HBSS (control), 10μM KU55933, 100mM KBrO_3_, or 10μM KU55933 and 100mM KBrO_3_ for 30 minutes. Quantification was completed with CellProfiler. For each condition, approximately 12 images were taken with 20-**40** cells per image. **b.** RPE1 cells were incubated with either 10μM scramble or ATM siRNA for 3**6** hours. Then cells were treated with either HBSS (control) or 100mM KBrO_3_ for 30 minutes. Quantification completed as described in a, with 15 images taken. **c.** SCA1 or wild type fibroblast cells were treated with either HBSS (control) or 100mM KBrO_3_ for one hour. Quantification completed as described in a, with 30 images taken. Potassium bromate treated conditions are both significantly different from controls (p ≤ 0.0001). **d.** Ataxin-1 interacts with ATM in response to oxidative stress. RPE1 cells were treated as described in a, then lysed in RIPA buffer. Lysates were incubated with the ATM antibody NB100-104 followed by protein G**-** sepharose beads. Immunoprecipitates were washed with RIPA buffer and interacting proteins were separated by SDS-PAGE and immunoblotted with the indicated antibodies. Image is representative of four or five independent experiments. One outlier removed by Grubb’s test.

Our next question was if reduction of total ATM protein levels was adequate to decreased pS186pS188 ataxin-1 signal. Similar to our kinase inhibition experiments, after 36 hours of siRNA ATM knockdown there was a diminished pS186pS188 ataxin-1 response to oxidative stress compared to control cells (Fig. 3b.) This indicates that both ATM phosphorylation capacity and protein levels impact the presence of pS186pS188 ataxin-1 in response to stress.

Next, we investigated the levels of pS186pS188 ataxin-1 in wild type versus SCA1 human patient-derived fibroblasts immortalized by hTERT, termed TruSCA1. This is a cell model system with the highest potential genetic accuracy to human SCA1 disease with a control line not affected by SCA1 and avoiding transformation that affects TP53 pathways. TruSCA1 line was derived from primary fibroblasts from a 29-year-old affected male with onset of olivopontocerebellar atrophy type I and spinocerebellar ataxia due to a CAG DNA expansion in *ATXN1* of 52 repeats on one allele. Control (TruHD-Q21Q18F) and SCA1 (TruSCA1-Q52Q29M) fibroblasts were treated with HBSS or 100mM KBrO_3_ for 30 minutes. There was no significant difference in pS186pS188 ataxin-1 immunofluorescent signal with or without the polyglutamine expansion (Fig. 3c). Similarly, pS186pS188 ataxin-1 levels increased significantly following KBrO_3_ treatment compared to control for both wild type and SCA1 fibroblasts (Fig. 3c). This indicates that human endogenous ataxin-1 pS186pS188 levels increase in response to oxidative stress, but this signaling is not affected by polyglutamine expansion.

We wanted to explore if ataxin-1 and ATM directly interacted within cells, and if this interaction was impacted by oxidative stress and phosphorylation capacity of ATM. To examine this, we conducted a co-immunoprecipitation of ATM under conditions of oxidative stress and ATM inhibition (Fig. 3d). Compared to control, there was a decrease in ataxin-1 in anti-ATM immunoprecipitated from cells treated with 100mM KBrO_3_ (Fig. 3d). However, if ATM phosphorylation was inhibited during an oxidative stress event, then more ataxin-1 was present (Fig. 3d). From this data, we hypothesize that ATM phosphorylation of ataxin-1 is acting like an on/off switch, with ataxin-1 requiring phosphorylation of serines 186 and 188 prior to localizing away from ATM to sites DNA damage. Further research is required to fully elucidate this mechanism.

Due to our data connecting ataxin-1 to ATM phosphorylation, we then assessed whether SCA1 patient fibroblasts were deficient in DNA damage markers. First, we examined γH2AX (histone family 2A variant), a marker of double stranded breaks, following treatment with bleomycin (25). There was no significant difference in γH2AX levels between wild type and SCA1 fibroblasts in either control or bleomycin treated conditions (Fig. 4a). Next we examined levels of 8-oxoguanine glycosylase (OGG1), an enzyme in the base excision repair pathway, after oxidative stress (26). Following KBrO_3_ treatment, SCA1 fibroblasts had an impaired response compared to wild type cells (Fig. 4b). This suggests that dysfunction with ataxin-1 in SCA1 cells may be involved in the base excision repair pathway. Further inquiry is required to identify if SCA1 cells have impaired responses with other member of the base excision repair pathway.

**Figure 4.**
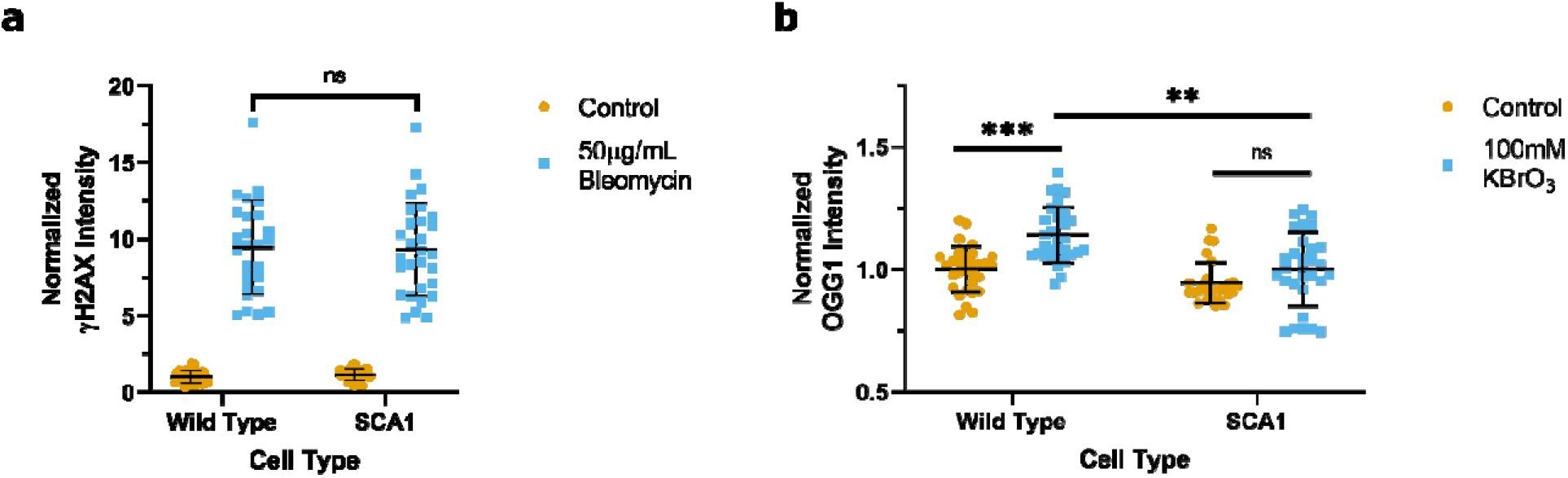
SCA1 cells have decreased oxidative stress response, similar double stranded break response to wild type. Quantification was completed with CellProfiler. Approximately 25-30 images were taken per condition with 20-40 cells per image. Significance values calculated by two-way ANOVA, ** = p ≤ 0.01, *** = p 0.001. Outliers identified by ROUT. **a.** Wild type and SCA1 fibroblasts were treated with either HBSS (control) or 50μg/mL bleomycin for 1 hour. Nuclear γH2AX signal was quantified. Bleomycin treated conditions are both significantly different from controls (p ≤ 0.0001) **b.** Wild type and SCA1 fibroblasts were treated with either HBSS (control) or 100mM KBrO_3_ for 30 min. Nuclear OGG1 signal was quantified.

**Figure 5.**
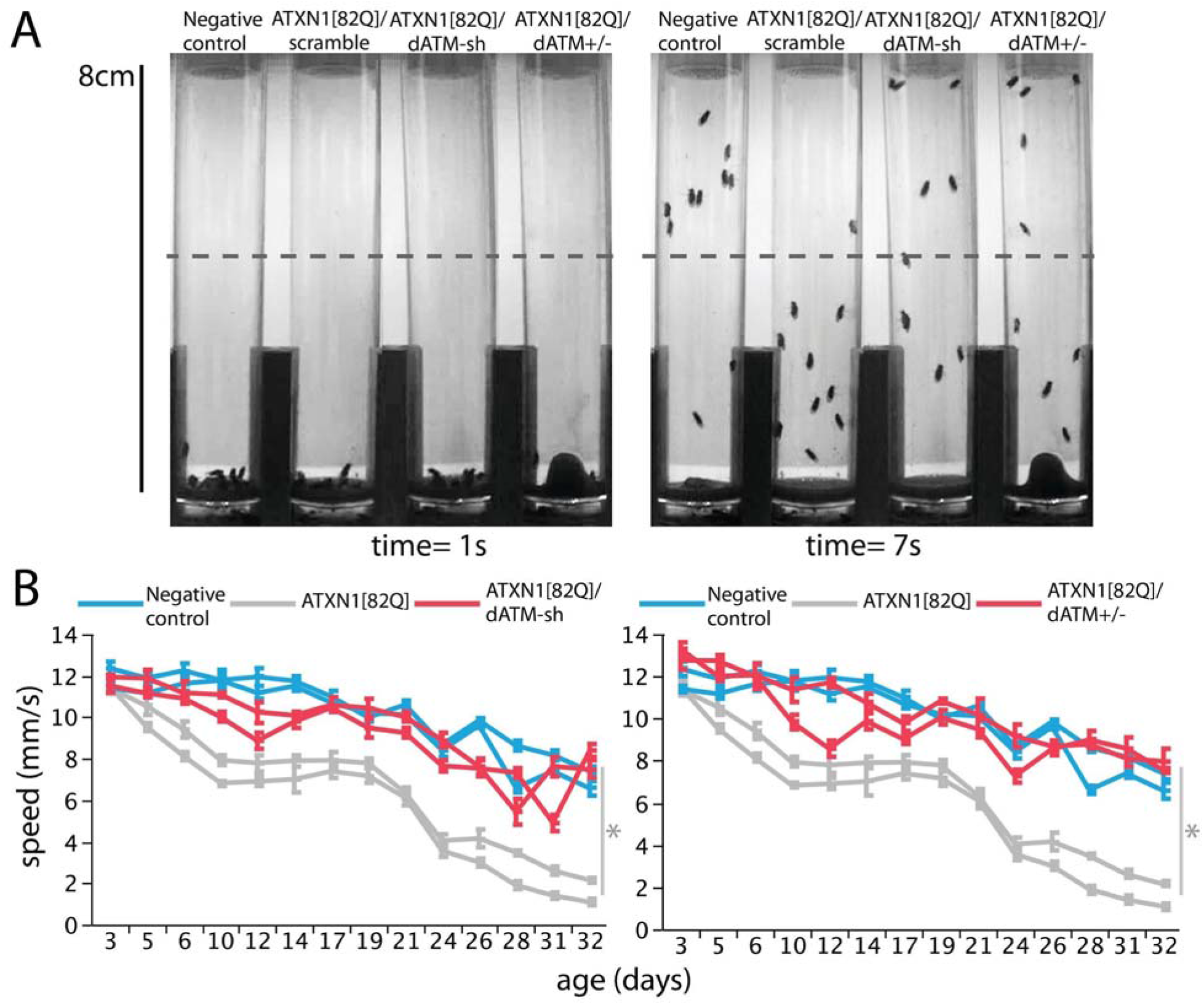
Knockdown of the *Drosophila* ATM homolog ameliorates ATXN1[82Q]-induced nervous system dysfunction. **a**, Time-sequence images illustrating the behavior of 24-day-old *Drosophila* females in a motor performance test. At 1s, all 10 age-matched virgin females of identical genotype were tapped to the bottom of a vial. After 7s, most of the negative control (healthy) fruit flies were able to climb past the midway point of the vial, whereas the majority of flies expressing ATXN1[82Q] in the nervous system (nerv-GAL4, also expressing a scramble, non-targeting shRNA) were at the bottom of the vial. ATXN1[82Q]-expressing animals in which the *ATM* fly homolog was knocked down specifically in the nervous system using an inducible shRNA (ATXN1[82Q]/dATM-sh), or carrying a heterozygous loss-of-function mutation in *dATM* (ATXN1[82Q]/dATM+/-), showed amelioration of ATXN1[82Q]-induced motor impairments. **b**, Graphs representing climbing speed as a function of age in healthy control fruit flies (blue lines), or in flies expressing ATXN1[82Q] alone (gray lines), or together with the dATM-sh or the dATM+/- loss-of-function allele (red lines). Two replicates are shown for each genotype. Error bars indicate s.e.m. The data was analyzed using linear mixed models ANOVA using a total of four independent replicates per genotype. *=p<0.0001 between ATXN1[82Q] and either ATXN1[82Q]/dATM-sh or ATXN1[82Q]/dATM+/-.

To validate our observations *in vivo*, and to assess the effect of reduced ATM function on ATXN1[82Q]-induced nervous system dysfunction, we used a well-characterized *Drosophila* SCA1 model^18^. Nervous system-specific expression of ATXN1[82Q] (using the *nrv2-GAL4* driver) leads to progressive motor performance deficits that can be quantified using movement metrics such as speed in a vial climbing assay. At 32 days of age, ATXN1[82Q] animals are 5 times slower than healthy controls in climbing assay (Fig. 3 a and b, compare grey lines with blue lines in b). Reduced function of the ATM *Drosophila* homolog, *tefu*, using either an inducible shRNA in the nervous system, or a heterozygous loss of function mutant, robustly ameliorates the ATXN1[82Q]-induced motor deficits (Fig. 3 a and b, compare grey lines with red lines in b). These data indicate that partial reduction of *tefu* levels protects from ATXN1[82Q]-induced spinocerebellar ataxia in a *Drosophila* model of SCA1.

## DISCUSSION

In this study, we describe the localization of ataxin-1 to sites of DNA damage and that this response is dependent on ATM kinase activity. Additionally, we discovered an ATM substrate site within ataxin-1 and provide evidence that ATM phosphorylates ataxin-1 at serine 188. Both ATM kinase inhibition and siRNA knockdown result in lower levels of pS186pS188 ataxin-1. We present preliminary evidence that ATM phosphorylation of ataxin-1 modulates ATM-ataxin-1 protein-protein interaction. We also demonstrate that wild type and SCA1 fibroblasts have similar levels of γH2AX, but decreased levels of OGG1 following stress treatment. Finally, we show that reducing levels of the *Drosophila* ATM homolog completely restores motor performance in a SCA1 fly model.

The data suggests that ATM signaling DNA damage repair via ataxin-1 is not affected by the SCA1 polyglutamine expansion. However, we observed a slower dynamic response of mutant ataxin-1 puncta to the regions of DNA suggesting impeded mutant ataxin-1 dynamics may result in poorer DNA damage repair. This is consistent with HD and SCA1 common modifier genes involved in DNA damage repair.

With the identification of the ataxin-1 response to DNA damage, SCA1 joins many other forms of ataxia with pathogenic mechanisms linked to DNA repair (27,28). This includes ataxia with ocular apraxia type 1 (AOA1), AOA2, Spinocerebellar ataxia with axonal neuropathy, and ataxia oculomotor apraxia XRCC1 (14). Dysfunction of ATM itself causes a form of recessive ataxia (29). More recently, the SCA3 protein ataxin-3 has been implicated in transcription coupled repair (30). Preliminary evidence has also suggested that alpha-synuclein can modulate DNA repair in Parkinson’s disease (31).

Overall, present findings that heterozygous loss-of-function of ATM homolog *tefu* or knockdown restores motor performance in a SCA1 fly model supports pursuing ATM kinase or protein expression inhibition as a potential therapeutic target for SCA1. Future work is needed to validate whether the best target is enzymatic modulation of ATM kinase, or lowering ATM protein levels, as this DNA repair factor is both an enzyme and a protein scaffold. This will lead to a better understanding of SCA1 pathology and open new avenues to developing potential therapeutics that may impact a broad range of genetic age-onset neurodegenerative diseases.

## MATERIALS AND METHODS

### Reagents

All reagents were sourced from Sigma-Aldrich, unless otherwise specified.

### Antibodies

The antibodies against phosphorylated serines 13 and 16 of the huntingtin N17 domain were previously characterised and validated(32). Anti-ataxin-1 antibodies were from Santa Cruz Biotechnology (sc-8766). Anti-rabbit IgG conjugated to Alexa Fluor 594 and anti-goat IgG conjugated to Alexa Fluor 488 were from ThermoFisher Scientific. For the phospho-specific antibody generation; antibody was raised in New Zealand white rabbits to NH_3_-G(p)SL(p)SQTPG-COOH, counter-purified over a non-phosphorylated peptide column then affinity purified over the phospho peptide column by a service from New England Peptides (MA, USA). See supplementary methods for details.

### Cells

Human retinal pigmented epithelial cells immortalized with hTERT (RPE1) were from the American Type Culture Collection. RPE1 cells were cultured in DMEM/F-12 1:1 media supplemented with 10% fetal bovine serum and 0.26% NaHCO_3_ at 37°C in a 5% CO_2_ incubator under nitrogen control of oxygen levels to 4%.

TruHD-Q21Q18F wild type cells were generated and cultured as described previously(33). TruSCA1-Q52Q29M hTERT immortalized fibroblasts were generated as follows: SCA1 patient fibroblasts were purchased from the Coriell Institute repository (GM06927). Cells were cultured in MEM with 15% fetal bovine serum and 1X GlutaMAX (Life Technologies #35050). Cells were infected with 1 × 10^6^ TERT Human Lentifect Purified Lentiviral Particles (GeneCopoeia, LPP-Q0450-Lv05-200-S). To aid in infection, 10 μg/ml polybrene was added. After 8 h, cells were infected again and left for 24 h. Media was changed, and cells were left for an additional 48 h. Successfully transduced cells were selected in media with 1 μg/ml puromycin. Cells were grown at 37°C with 5% CO_2_ and 4% oxygen.

### Transfections

All cells were transfected with TransIT-X2 (Mirus Bio) according to the manufacturer’s specifications. All transfections involving ataxin-1 constructs were incubated at 37°C for 8 hours with 1 μg of DNA. Ataxin-1 expression plasmids; eGFP-ataxin-1 [Q26], eGFP-ataxin-1 [Q84]; were generated as described previously(17). EGFP-ataxin-1 [Q26] S186A and eGFP-ataxin-1 [Q26] S188A were generated from eGFP-ataxin-1 [Q26] using Q5 Site-Directed Mutagenesis Kit according to the manufacturer’s specifications. All PCR reagents and enzymes were purchased from New England Biolabs. Plasmids were purified via Presto Mini Plasmid Kit (Geneaid) and sequences were verified by PCR sequencing by the McMaster Mobix facility.

### Immunofluorescence

Cells were fixed using methanol at −20°C for 20 minutes, then washed with wash buffer (50mM Tris-HCl, pH 7.5, 150mM NaCl, 0.1% Triton X-100). Then, cells were blocked with blocking buffer (wash buffer + 5% FBS) for either 10 minutes at room temperature or overnight at 4°C. Cells were incubated with primary antibodies diluted in blocking buffer for 1 hour at room temperature. Next, cells were washed and then incubated with secondary antibodies diluted in blocking buffer for 20 minutes at room temperature. Following washing with wash buffer, cells were imaged in PBS.

### Microscopy

All microscopy was completed using a Nikon A1 confocal system attached to a Nikon Eclipse Ti inverted microscope, using an PLAN APO 60x/1.40 oil objective or PLAN APO 20×/0.75□dry objective with Spectra X LED lamp (Lumencor) and GaAsP detectors. A 405nm laser which was part of the Nikon A1 confocal system was used for irradiation experiments. 405nm, 489nm, and 561nm lasers experiments were used for imaging.

### Micro-irradiation Assay

RPE1 cells were grown in glass-bottom six well tissue culture dishes or eight well μ-slide ibiTreat until 80-85% confluence, then stained with NucBlue (ThermoFisher Scientific) for 15 minutes at 37°C in a 5% CO_2_ incubator. Media was aspirated and replaced with Hank’s Balanced Salt Solution (HBSS) (ThermoFisher Scientific) immediately preceding irradiation. Using the Nikon A1 confocal setup described in the subsequent section, samples were kept at 37°C with a Tokai Hit Inu Incubation system (model WSKM). A 405nm laser set to 100% power was used to irradiate regions of interest using a scan speed of 1/16 frames per second (512 x 512 pixels). Regions of interest were drawn over areas where ataxin-1 nuclear inclusions were absent. After X-Y coordinates were recorded, cells were either imaged live, or incubated at 37°C for specified incubation times preceding methanol fixations and immunofluorescence. Then X-Y coordinates were revisited for imaging. For live cell imaging, cells transfected with ataxin-1 constructs were irradiated as described above. Cells were imaged at either 1 minute, 2 minute, or 5 minute intervals over a thirty-minute period using confocal microscopy as described in the subsequent section. For ATM kinase inhibition trials, cells were incubated for 15 minutes in HBSS with 10μM ATM kinase inhibitor KU55933 prior to irradiation.

### Average Pixel Intensity Quantification

Average pixel intensity of the regions of interest were measured using ImageJ mean gray function on FITC channel image. Higher intensity indicates presence of ataxin-1 nuclear inclusions. Values were normalized to the average of pre-irradiation control cell values, which had a region of interest defined but were not irradiated. An increase in intensity indicates that fluorescent puncta are localizing into the region of interest. This analysis was conducted at three time points: 0 minutes, 10 minutes, 25 minutes, and 40 minutes. Statistical analysis, including Student’s t-test and ANOVA, were completed using GraphPad Prism software.

### Chromatin Retention Assay

RPE1 cells were transfected with indicated eGFP-ataxin-1 construct and transfection control histone 2B-mCherry (H2B-mCherry). After 24 hours incubation at 37°C, cells treated with either HBSS (control) or 100mM KBrO_3_ in HBSS for 30 minutes. Soluble proteins were extracted with cold 0.2% Triton X-100 in PBS for 2 minutes on ice, then cells were fixed with 4% paraformaldehyde for 15 minutes at room temperature. Nuclear intensity was quantified using CellProfiler(34), using H2B mCherry to normalize for variability in transfection efficiency. Intensity was normalized to untreated wild type ataxin-1 conditions.

### pS186pS188 Ataxin-1 Oxidative Stress Trials

For trials focusing on ATM phosphorylation capacity, RPE1 cells were incubated at 37°C for 30 minutes with either HBSS (control), 10μM KU55933 in HBSS, 100mM KBrO_3_ in HBSS, or 10μM KU55933 and 100mM KBrO_3_ in HBSS. For trials focusing on ATM protein levels, endogenous ATM knockdown was established with 10 μM ATM SMARTpool siGENOME siRNA (Dharmacon, M003201-04-0005) in RPE1 cells. Control cells were treated with 10μM scramble siRNA (Santa Cruz, sc-37007). siRNA was transfected with Lipofectamine RNAiMax (Invitrogen) according to the manufacturer’s instructions, then incubated for 36 hours at 37°C. Cells were treated with either HBSS (control) or 100mM KBrO_3_ in HBSS for 30 minutes. Cells were then fixed with methanol and immunofluorescence was conducted as described above. Raw nuclear intensity was quantified using CellProfiler, with values being normalized to the average of the HBSS control values.

### Co-Immunoprecipitation

RPE1 cells were incubated at 37°C for 30 minutes with either HBSS (control), 10μM KU55933 in HBSS, 100mM KBrO_3_ in HBSS, or 10μM KU55933 and 100mM KBrO_3_ in HBSS. Cells were trypsinized, resuspended in PBS, then fixed with 1% PFA for 10 minutes. The fixation reaction was quenched with 1M glycine, then lysed in RIPA buffer (50□mM Tris-HCl pH 8.0, 150□mM NaCl, 1% NP-40, 0.25% sodium deoxycholate, 1□mM EDTA, protease and phosphatase inhibitors (Roche)). Input samples were acquired, then remaining lysates were incubated anti-ATM (Novus Biologicals, NB100-104) and protein A sepharose beads overnight with rotation at 4°C. Beads and proteins were washed three times with RIPA lysis buffer, then denatured with SDS-loading buffer at 100°C for 10□minutes. Samples were then separated by SDS-PAGE and analyzed by western blot with anti-ATM (sc-377293) and anti-ataxin-1 (11750, kind gift from Zoghbi Laboratory (35)) as outlined in supplemental methods.

### *Drosophila* Models and Motor Performance Tests

The *Drosophila* SCA1 model and transgenic lines expressing human ATXN1[82Q] were previously described (36). The inducible shRNA line v108074 specifically targeting *Tefu*, the *Drosophila ATM* orthologue, and the control, non-targeting, scramble shRNA v2691 were obtained from the Vienna Drosophila Stock Center. The nervous system driver line *nrv2-GAL4*, and the *Tefu (dATM)* loss-of-function allele *Mi{ET1}tefuMB09945* were obtained from the Bloomington Drosophila Stock Center. For the *Drosophila* motor performance assay, we used an automated data acquisition system. This system taps the vials at 7s intervals and records video files. These files are then processed using a custom-built software that calculates the average speed of the animals in each vial. 10 age-matched virgin females were used per replicate, and at least four replicates per genotype were tested. Animals are transferred into vials containing fresh media daily. For statistical analysis we performed linear mixed models ANOVA between the indicated genotypes using four replicates per genotype. All cultures, breeding and tests were performed at 25°C.

### *Drosophila* Models Genotypes

<u>negative controls</u>: w[1118]/+, nrv2-GAL4/UAS-v2691.
<u>ATXN1[82Q]/scramble</u>: UAS-ATXN1[82Q]F7/w[1118]; nrv2-GAL4/UAS-v2691.
<u>ATXN1[82Q]/dATM-sh1</u>: UAS-ATXN1[82Q]F7/w[1118]; nrv2-GAL4/UAS-v108074.
<u>ATXN1[82Q]/dATM+/-</u>: UAS-ATXN1[82Q]F7/w[1118]; nrv2-GAL4/+;
Mi{ET1}tefuMB09945/+

## ACKNOWLEDGMENTS

We thank H. Zoghbi (Baylor College of Medicine) for her kind gift of ataxin-1 11750 antibody.

This work is supported by the Krembil Foundation and the Canadian Institutes for Health Research (CIHR) Frederick Banting and Charles Best Canada Graduate Scholarships Doctoral Awards (CGS D), Grant/Award Number: FBD-170797.

## CONFLICT OF INTEREST STATEMENT

None declared.

## ABBREVIATIONS

AOA1: Ataxia with ocular apraxia type 1
AOA2: Ataxia with ocular apraxia type 2
ATM: Ataxi a-tel angi ectasi a mutated
HBSS: Hank’s balanced salt solution
HD: Huntington’s Disease
OGG1: 8-Oxoguanine glycosylase
RPE1: Retinal pigment epithelial cells
SCA1: Spinocerebellar ataxia type 1
SCA3: Spinocerebellar ataxia type 3
SCA7: Spinocerebellar ataxia type 7
γH2AX: Histone family 2A variant

## Notes

### Competing Interest Statement

The authors have declared no competing interest.

### Summary of Updates

Some expanded text, data added to figures 2 and 3.

