## Supplemental data for "Spinocerebellar Ataxia Type 1 protein Ataxin-1 is signalled to DNA damage by Ataxia Telangiectasia Mutated kinase"

### SUPPLEMENTARY MATERIAL

#### ***Phospho-Specific Antibody Generation***

The antibody was raised in New Zealand white rabbits to NH_3_-G(p)SL(p)SQTPG-COOH, then affinity-purified over a peptide column. All steps, including peptide synthesis, injections, titrations, bleeds, and affinity purification, were carried out at New England Peptide (Garner, MA, USA) with the standard PSSA Rabbit package. The antibody was then validated by dot-blot assay to determine epitope specificity; western blotting following siRNA knockdown to determine protein specificity; and peptide competition assay to determine immunofluorescent signal specificity.

*Dot Blot Assay*

Serial dilution concentrations (25, 50, 100, 500, 1000ng) of synthetic ataxin-1 peptides (non-phospho GSLSQTPG, phospho GpSLpSQTPG, and GALAGTPG) were spotted onto a nitrocellulose membrane (Pall Life Sciences) and were dried at room temperature for 45 minutes.

The membrane was blocked with 5% non-fat milk powder in TBS-T (50 mM Tris-HCl, pH 7.5, 150 mM NaCl, 0.1% Tween-20) for 1 hour at room temperature. Amido black protein staining was conducted as outlined in Goldman et al. and imaged (1). The membrane was then washed in TBS-T for 1 hour at room temperature, then incubated with primary phospho-ataxin-1 antibody (1:1000) overnight at 4°C. The membrane was washed for 10 minutes three times with TBS-T, then incubated with anti-rabbit peroxidase (HRP) secondary (1:50,000; Abcam) for 45 minutes at room temperature. The membrane was again washed for 10 minutes three times with TBS-T, then imaged with enhanced chemiluminescent HRP substrate (EMD Millipore) on a MicroChemi system (DNR Bio-imaging Systems).

*siRNA Knockdown*

Endogenous ataxin-1 knockdown was established with ataxin-1 SMARTpool siGENOME siRNA (Dharmacon, M-004510-02-0005) in RPE1 cells. siRNA was transfected with Lipofectamine RNAiMax (Invitrogen) according to the manufacturer’s instructions. Control dishes were established with scrambled siRNA.

Cells were lysed in radioimmunoprecipitation assay buffer with 10% phosphatase (Roche) and 10% protease inhibitors (Roche), scrapped, and incubated on ice for 15 minutes. Cell lysates were centrifuged at full speed for 10 minutes at 4°C. Protein concentration was quantified by Bradford assay, then 40-60 µg of protein was loaded into a precast 4–20% polyacrylamide gradient gel (Biorad). Proteins were then separated by SDS–PAGE and electroblotted onto 0.45 µm polyvinyl difluoride (PVDF) membrane (EMD Millipore). Western immunoblotting was then conducted.

*Western Immunoblotting*

Immunoblots were blocked with 5% non-fat milk powder in TBS-T (50 mM Tris-HCl, pH 7.5, 150 mM NaCl, 0.1% Tween-20) for 1 hour at room temperature. Blots were cut horizontally at a 48-kDa marker to probe for ataxin-1 (87 kDA) or GAPDH (loading control, 37 kDa) separately. Blots were then incubated with primary phospho-ataxin-1 antibody (1:1000) or GAPDH (1:10,000; Abcam ab8425) overnight at 4°C. Blots were washed for 10 minutes three times with TBS-T, then incubated with anti-rabbit peroxidase (HRP) secondary (1:50,000; Abcam) for 45 minutes at room temperature. Blots were again washed for 10 minutes three times with TBS-T, then imaged with enhanced chemiluminescent HRP substrate (EMD Millipore) on a MicroChemi system (DNR Bio-imaging Systems). Bands were quantified using National Institutes of Health ImageJ. Phospho-ataxin-1 signal was normalized to the GAPDH loading control.

*Immunofluorescence Peptide Competition Assay*

The phospho-ataxin-1 antibody (1:250 was incubated with 1000 ng of synthetic peptides (non-phospho ataxin-1, phospho ataxin-1, and p53 control peptide) at room temperature for 1 hour. RPE1 cells were then fixed with cold methanol as previously described, blocked with 10% FBS in TBS +0.1% TritX for 1 hour, then incubated overnight with the treated antibodies. Cells were washed three times with PBS, then incubated with anti-rabbit AlexaFluor488 secondary antibody (1:500, Molecular Probes) for 45 minutes at room temperature. Cells were then washed three times with PBS, then imaged as previously described.

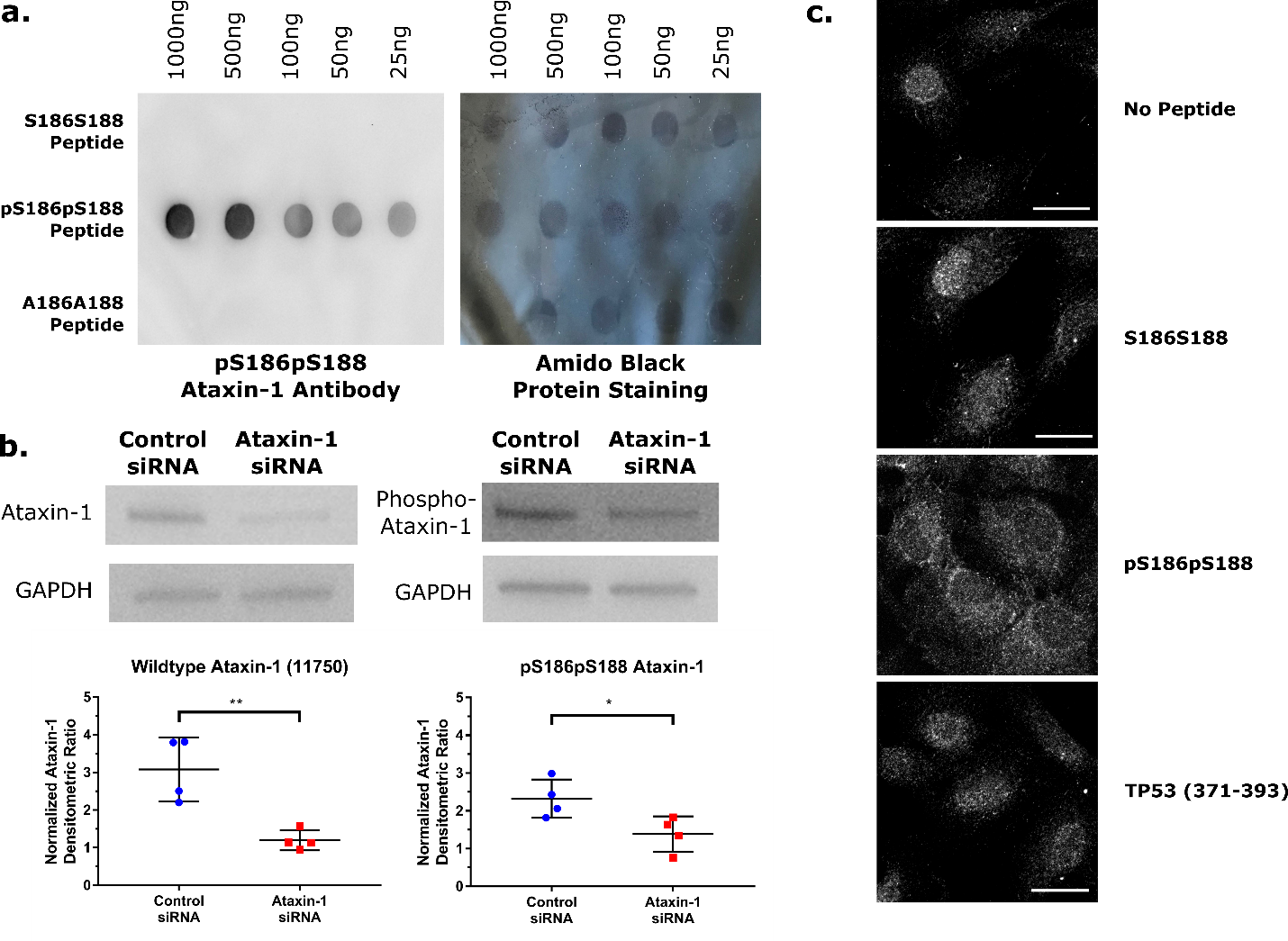

**Figure S1. pS186pS188 ataxin-1 antibody validation. a.** Dot blot assay with ataxin-1 peptide in the unphosphorylated (S186S188), phosphorylated (pS186pS188), and alanine substitution state (A186A188). Protein staining was conducted with 1X Amido Black. **b.** Ataxin-1 knockdown showing pS186pS188 antibody specificity compared to previously validated 11750 antibody. Blots were cut at the 48 kDa marker to probe for ataxin-1 and GAPDH separately. p= 0.0063 for 11750 ataxin-1, p= 0.0767 for pS186pS188 ataxin-1. Analyzed by unpaired t-test of three independent replicates. Error bars indicate standard deviation. **c.** Peptide competition assay with unphosphorylated S186S188, phosphorylated pS186pS188, and non-specific TP53 peptide. The cytoplasmic signal is non-specific. Note that in no peptide control, S186S188, and TP53 there is an evenly distributed signal in the nucleus, with transient stress events causing an increase in nuclear signal for some cells. In pS186pS188, the nuclear signal is reduced and no increase in nuclear signal is observed.

**Video S1. Live Cell Imaging eGFP-ataxin-1 [Q26] with 10µM KU55933.** Laser microirradiation assay of RPE1 cells transfected with eGFP-ataxin-1[Q26] after incubation with 10μM ATM kinase inhibitor KU55933 imaged at 1 frame per minute.
